# Category-biased patches encircle core domain-general regions in the human lateral prefrontal cortex

**DOI:** 10.1101/2025.01.16.633461

**Authors:** Moataz Assem, Sneha Shashidhara, Matthew Glasser, John Duncan

## Abstract

The fine-grained functional organization of the human lateral prefrontal cortex (PFC) remains poorly understood. Previous fMRI studies delineated focal domain-general, or multiple-demand (MD), PFC areas that co-activate during diverse cognitively demanding tasks. While there is some evidence for category-selective (face and scene) patches, in human and non-human primate PFC, these have not been systematically assessed. Recent precision fMRI studies have also revealed sensory-biased PFC patches adjacent to MD regions. To investigate if this topographic arrangement extends to other domains, we analysed two independent fMRI datasets (n=449 and n=37) utilizing the high-resolution multimodal MRI approaches of the Human Connectome Project (HCP). Both datasets included cognitive control tasks and stimuli spanning different categories: faces, places, tools and body parts. Contrasting each stimulus category against the remaining ones revealed focal interdigitated patches of activity located adjacent to core MD regions. The face and place results were robust, replicating across different executive tasks, experimental designs (block and event-related) and at the single subject level. In one dataset, where participants performed both category and sensory tasks, place patches overlapped with visually biased regions, while face patches were positioned between visual and auditory biases. Our results paint a refined view of the fine-grained functional organization of the PFC, revealing a recurring motif of interdigitated domain-specific and domain-general circuits. This organization offers new constraints for models of cognitive control, cortical specialization and development.

## Introduction

The primate prefrontal (PFC), in coordination with wider brain networks, plays a critical role in integrating diverse neural signals in support of complex cognitive operations (Banich et al., 2024; Miller and Cohen, 2001). A central open question is the extent to which the PFC is organized into functionally specialized modules versus domain-general processes. Human functional magnetic resonance imaging (fMRI) studies converge on a circumscribed set of domain-general or multiple-demand (MD) regions that co-activate during a diverse set of multiple cognitive demands (Assem et al., 2020; Cole and Schneider, 2007; Duncan, 2010; Fedorenko et al., 2013). We recently used the high-resolution multimodal imaging approaches of the Human Connectome Project (HCP) to delineate 9 MD cortical patches, 5 within the lateral PFC (Assem et al., 2024, Assem et al., 2020) (**Figure 1a**). MD regions’ fine-grained activation patterns flexibly represent behaviourally relevant stimuli and task rules (Woolgar et al., 2016). Single-neuron recordings in putatively homologous MD regions in non-human primates (NHPs) respond to diverse mixtures of behaviourally-relevant signals (Rigotti et al., 2013; Stokes et al., 2013).

**Figure 1.**
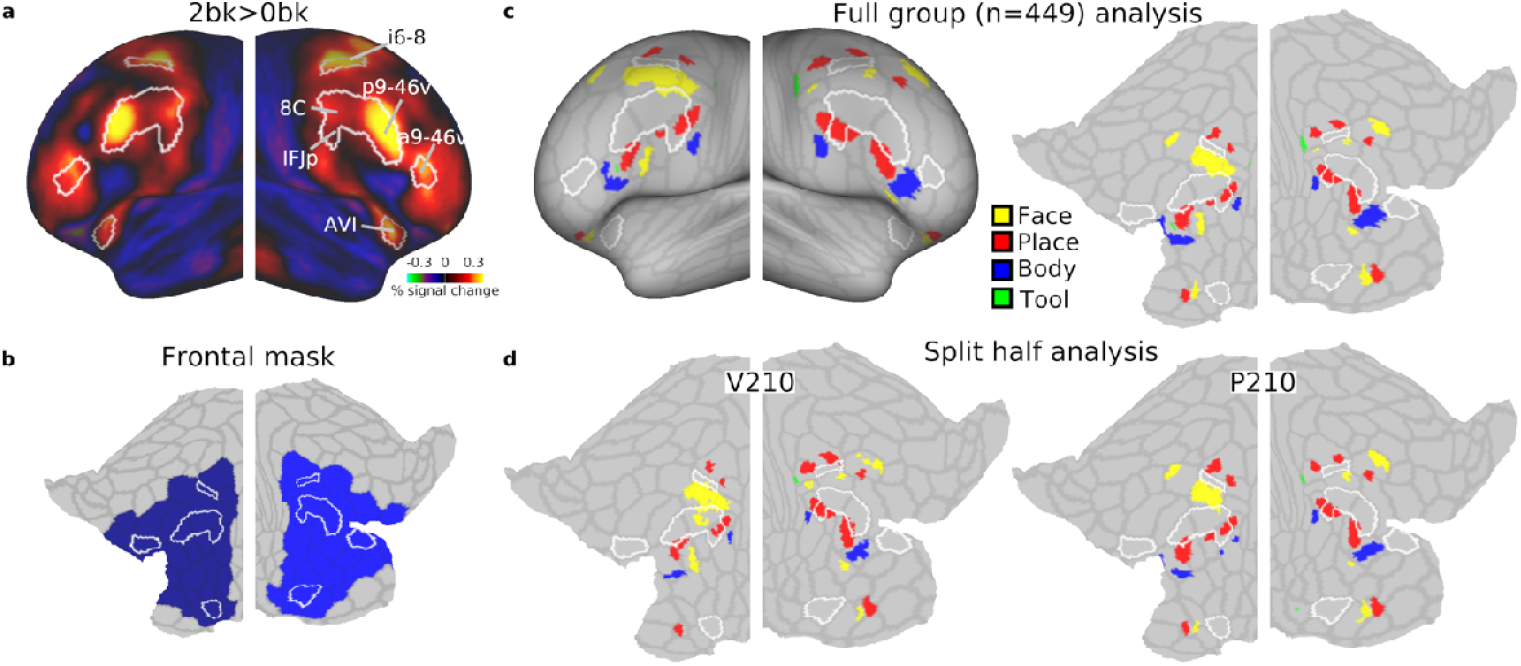
Category-biased patches in the HCP dataset. (a) Group average activation map for the 2bk>0bk contrast displayed on bilateral inflated cortical surface. Core MD regions (white borders) are defined based on (Assem et al., 2020). **(b)** Flat map displaying pre-defined bilateral frontal mask used for statistical analysis. It encompasses core MD and their adjacent areas. The HCP MMP1.0 borders are displayed in faint gray here and in other flat maps throughout the manuscript. Note that non-lateral frontal core MD areas are not displayed on any flat map. **(c)** Statistically defined category-biased clusters displayed on inflated (left) and flattened (right) cortical surfaces. **(d)** Split-half analysis showing category-biased clusters for two-subgroups, demonstrating the replicability of findings. Data available at https://balsa.wustl.edu/V8XqK.

At the same time, there is evidence for domain-specificity within circumscribed regions of the primate PFC. Studies in NHPs have identified neurons tuned to specific categories, such as faces (Ó Scalaidhe et al., 1997; Rolls et al., 2006; Romanski, 2012; Romanski and Diehl, 2011) and objects (Rozzi et al., 2021) within localized PFC regions. FMRI findings corroborate this, showing distinct face and scene or place patches mainly in the ventral frontal cortex of NHPs (Haile et al., 2019; Li et al., 2022; Schaeffer et al., 2020; Tsao et al., 2008) and humans (Chan and Downing, 2011; Henderson et al., 2020; Rajimehr et al., 2009; Troiani et al., 2016). However, the relationship between these domain-specific patches and MD regions remains poorly understood.

Recent work using precision fMRI approaches revealed a side-by-side spatial arrangement of sensory-biased areas, tuned to auditory or visual inputs, with frontal MD regions (Assem et al., 2022; Michalka et al., 2015; Noyce et al., 2022; Tobyne et al., 2017) [see also (Fedorenko et al., 2012)]. This interdigitated organization, we hypothesised, might facilitate the integration of specialized sensory signals into a domain-general workspace (Assem et al., 2022; Duncan et al., 2020). Here we investigate whether a similar arrangement exists for category-selective areas, such as those processing faces and places. A previous study using multivariate analysis suggest that regions showing strong decoding of faces vs places lie adjacent and ventral to frontal MD regions (Shashidhara et al., 2024).

To explore this, we analysed two independent datasets: the young adult HCP dataset (Barch et al., 2013; Glasser et al., 2016), and the Cognition and Brain Sciences Unit (CBU) dataset (Assem et al., 2024, 2022), which also used HCP-style acquisition and analysis approaches (Glasser et al., 2016). These datasets included 5 diverse tasks featuring visual stimuli, including faces, places, tools and body parts. Critically, both datasets also included task difficulty manipulations, known to characteristically engage MD regions. Our results reveal an expanded set of focal category-biased patches in the PFC. Importantly, these patches predominantly spare but lie adjacent to MD territories. These findings point to a highly structured and fine-grained organization within the PFC, where interdigitated domain-specific and domain-general patches may represent a fundamental organizational principle, facilitating the integration of signals from functionally segregated systems.

## Methods

### Participants

#### HCP dataset

The analyzed dataset consisted of 449 healthy volunteers from the HCP S500 release. Subjects were recruited from the Missouri Twin Registry (186 males, 263 females), with age ranges 22–25 (n□=□69), 26–30 (n□=□208), 31–35 (n□=□169), and 36+ (n□=□3). Informed consent was obtained from each subject as approved by the institutional Review Board at Washington University in St. Louis.

#### CBU dataset

Data from 37 human participants were included in this study (age = 25.9□±□4.7, 23 females, all right-handed). Informed consent was obtained from each subject and the study was approved by the Cambridge Psychology Research Ethics Committee.

### Task paradigms

#### HCP n-back task

Task details are detailed in (Barch et al., 2013). Briefly, each run consisted of eight task blocks (10 trials of 2.5 s each, for 25 s) and four fixation blocks (15 s each). Within each run, four blocks used a 2-back task (respond “target” whenever the current stimulus was the same as the one 2-back) and the other four used a 0-back task (a target cue was presented at the start of each block, and a “target” response was required to any presentation of that stimulus during the block). Stimuli consisted of pictures of faces, places, tools, and body parts; within each run, the four different stimulus types were presented in separate blocks.

#### CBU executive function (EF) tasks

Details of the task paradigms are reported in (Assem et al., 2024). Participants performed three visual tasks—n-back, switch, and stop signal—designed to probe executive functions while engaging with face and place stimuli. Each trial featured a visual stimulus displayed for 1500 ms, followed by a 500 ms blank screen. Stimuli were presented in separate blocks by category, with faces depicting males or females, children or adults, with happy or sad expressions (from the Developmental Emotional Faces Stimulus Set, Meuwissen et al., 2017), and places including images of houses or churches, old or new, from interior or exterior views. There were 32 unique stimuli per category, evenly distributed across 2 × 2 × 2 feature combinations.

The *N-back task* involved monitoring a sequence of stimuli to identify repetitions: for the 1-back condition (easy), participants responded to immediate repeats, while for the 3-back condition (hard), they responded to repetitions three trials back, with 1–2 targets and 2 lures per block. An auditory version of the n-back task was administered to the same participants in a separate session, with identical design to the visual version. Auditory stimuli included 9 animate (e.g. a human’s cough, a lion’s roar) and 9 inanimate (e.g. a musical instrument, a bell ringing) sounds. For further details, see (Assem et al., 2022).

The *Switch task* required responses based on two rule types indicated by colored borders (red or blue). In easy blocks, a single rule applied throughout (e.g., classify faces by gender or age, or places by type or setting). In hard blocks, the border color switched randomly, requiring participants to alternate between rules.

The *Stop signal task* involved categorizing stimuli (happy vs. sad faces, old vs. new places) in no-stop blocks, while in stop blocks, 33% of trials included a stop signal (a black circle), instructing participants to withhold their response. Stop signal delays (SSD) were adjusted dynamically based on reaction times to maintain task difficulty.

The *Temporal sequence task* was event-related and conducted during a second scanning session with the same participants who performed the EF task. It used the same face and place images as the EF tasks. Each trial involved presenting a sequence of three images at the center of the screen, which participants were instructed to memorise. In the execution phase, the same three images were presented again simultaneously on the screen. Participants used a keypad to press the sequence corresponding to the original presentation order. For example, if the images on the screen from left to right were the 3rd, 1st, and 2nd in the sequence, the correct answer would be: 3, 1, 2. During scanning, participants completed four runs of 48 trials each, resulting in a total of 192 trials. Each run included six repetitions of eight unique sequences (FFF, FFP, FPF, FPP, PFF, PFP, PPF, PPP; F stands for face, P for place), generated by all possible combinations of the two categories. For each sequence, random face or building images were assigned according to the category, with the restriction that no image was repeated within a sequence. Each trial began with a fixation cross displayed for 1 s, followed by a sequence of three images. Each image was displayed for 1 s, separated by a 1 s blank screen. In two-thirds of the trials (32 trials per run), the execution phase followed a 1 s delay. The trial concluded with a 1s inter-trial interval (blank screen). For the remaining one-third of the trials (16 trials per run), the execution phase was omitted, and the trial moved directly to the inter-trial interval. There were six possible response sequences, which were pseudo-randomized across trials. Each sequence appeared five times per run, with two additional trials containing random sequences.

### Image Acquisition

MRI acquisition protocols have been previously described (Assem et al., 2024, 2022; Glasser et al., 2013; Smith et al., 2013; U□urbil et al., 2013). For the HCP dataset, imaging data were acquired using a customized 3T Siemens “Connectom” scanner with a 100 mT/m SC72 gradient insert and a 32-channel RF receive head coil. Structural scans included at least one T1-weighted (T1w) MPRAGE and one 3D T2-weighted (T2w) SPACE scan at 0.7-mm isotropic resolution. Functional MRI (fMRI) data consisted of four 15-minute runs of resting-state (rfMRI) and seven task-based fMRI tasks totaling 46.6 minutes, all acquired using multi-band echo-planar imaging (EPI) sequences (TR = 720 ms, 2-mm isotropic resolution). Spin echo phase-reversed images with matched geometry to the fMRI were collected during fMRI sessions.

The CBU dataset utilized a 3T Siemens Prisma scanner with a 32-channel RF receive head coil, employing HCP Lifespan protocols. Each subject attended 2 scanning sessions. Structural scans included one T1w MPRAGE and one 3D T2w SPACE scan at 0.8-mm isotropic resolution. Functional data included two 15-minute rfMRI runs and four runs of five task-based fMRI tasks (∼100 minutes total) acquired with multi-band EPI sequences (TR = 800 ms, TE = 37 ms, 2-mm isotropic resolution). Both rest and task EPI runs were acquired in pairs of reversed phase-encoding directions (AP/PA). Spin echo phase-reversed images were also acquired during structural and functional scans in AP/PA directions with matched geometry to the fMRI.

### Data Preprocessing

The preprocessing for both datasets followed the HCP minimal preprocessing pipelines (Glasser et al., 2013) with some dataset-specific adjustments described previously(Assem et al., 2024; Glasser et al., 2016) and briefly outlined below. Structural images (T1w and T2w) were used to extract cortical surfaces and segment subcortical structures, while functional images (rest and task) were mapped from volume to surface space and combined with subcortical data to form the standard CIFTI grayordinate space. Surface based spatial smoothing was applied using a 2-mm full-width at half-maximum (FWHM) kernel. ICA□+□FIX (Salimi-Khorshidi et al., 2014) was applied identically to rest and task fMRI data.

Both datasets employed multimodal surface matching (MSMAll) for cross-subject cortical alignment, using cortical folding maps (“sulc”), myelin maps, and resting-state visuotopic maps (Robinson et al., 2018, 2014). Traditional fMRI alignment methods typically rely heavily on cortical curvature alone, which varies substantially across individuals - even in twins - and particularly in association cortices, leading to blurring across functional boundaries and reduced statistical power (Glasser et al., 2016). MSM works by describing the cortex as a 2-dimensional sheet or a sphere. This representation simplifies the co-registration process, making it computationally more manageable than prevailing 3D methodologies. MSMAll enhances alignment by first initializing registration based on prominent cortical folding patterns and then refining it using myelin and connectivity features, which are more consistently expressed across individuals. As a result, MSMAll achieves substantially better cross-subject alignment: the fraction of the individually defined regions that are captured by group-defined borders reaches 60%–70% (Coalson et al., 2018).

Key differences between preprocessing of the two datasets include the use of HCP pipeline version 4.0.0 for the CBU dataset compared to earlier versions for the HCP dataset. For the CBU dataset, an improved ICA□+□FIX classifier(Glasser et al., 2019, 2018) was used for task fMRI noise removal, and thresholds were optimized based on manual inspection of a subset of subjects. Unlike the HCP dataset, global structured noise (e.g., respiration-related noise) was not removed from the CBU dataset due to unavailable temporal ICA scripts at the time the dataset was preprocessed.

### General Linear Models of task fMRI data

Task fMRI analyses for both datasets followed standard procedures outlined in the previous publications (Assem et al., 2024; Barch et al., 2013) and briefly outlined below. For both datasets, activation estimates were computed for the preprocessed time series from each run using a general linear model (GLM) implemented in FILM. All regressors were convolved with a canonical hemodynamic response function and its temporal derivative. Fixed-effects analyses using FSL’s FEAT were conducted to estimate within-subject averages across runs. Beta “cope” maps were exported to MATLAB, corrected for intensity scaling, and converted to percent signal change (100 × [beta/10,000]).

For the HCP dataset, eight regressors were used for the working memory (WM) task (one for each stimulus type in the 0-back and 2-back conditions), with activation estimates derived from contrasts of interest (e.g., WM 2bk > 0bk).

For the CBU EF tasks, four regressors (stimulus category × task difficulty) were modelled for each task, with predictors spanning from the cue onset to the final trial offset (28 s) and including 12 motion regressors (translations, rotations, and their derivatives).

For the CBU temporal sequence task, separate regressors modelled the sequence display and execution phases, along with 12 motion regressors. The sequence display phase included eight regressors, one for each unique sequence category, covering the 6-second period from stimulus onset to the start of the delay. The execution phase was modelled with trial-wise regressors spanning from execution stimulus onset to trial end. From the display phase, we generated six ‘cope’ maps: Step1 Face, Step1 Place, Step2 Face, Step2 Place, Step3 Face, and Step3 Place. Each cope map was derived by averaging across the relevant regressors. For example, Step1 Face cope was created with a 0.25 weight of each of the 4 regressors where step1 was a face and all other regressors were 0. We then averaged across all steps to create a face map and a place map, which we directly contrasted to examine category-specific activations.

### Regions of interest (ROIs) definition and further fMRI analysis

The HCP multimodal parcellation (MMP1.0) divides the cortex into 180 areas. We have previously identified that the 9 MD patches overlap with 10 areas that we have labelled core MD as they showed the strongest co-activations and resting-state fMRI interconnectedeness (Assem et al., 2020). The lateral frontal cortex encompasses 6 core MD areas: a9-46v, p9-46v, 8C, IFJp, i6-8, AVI (white borders in Figure 1a).

The lateral PFC mask was defined to encompass lateral frontal core MD areas and any area that shared a border with them. Our reasoning was to capture category preferences in the immediate vicinity of core MD regions. Additionally, areas 44 and 45 were included to avoid gaps in the mask (Figure 1c).

For cluster definition we used the function “wb_command -cifti-find-clusters” from Connectome Workbench (v1.5.0) with subject-specific vertex areas and the midthickness cortical surface as inputs. We applied the function within the lateral PFC mask on the beta activation maps and not statistical maps (e.g. t-or z-stat map) as activation peaks are more directly linked to the true spatial locations and magnitude of neurobiological responses (Glasser et al., 2016). We did not specify any activation thresholds. Instead, we identified all clusters that were at least 5 mm^2^ in surface area and no smaller than 10% the size of the largest cluster, minimizing the definition of minute speckles as clusters. For further statistical analysis, activations of vertices sharing the same cluster label were averaged together to obtain a single value for each area.

We calculated gradient maps for the whole cortex using the function wb_command –cifti-gradient from Connectome Workbench (v1.5.0) with no presmoothing and using subject-specific vertex areas and the midthickness cortical surface as inputs.

## Results

### Mapping category-biased patches near core MD regions in the HCP dataset

We first analyzed the HCP n-back (2bk and 0bk) task which used stimuli from four categories: faces, places, tools and body parts. A hard vs easy n-back contrast is well established to strongly activate MD regions, and we previously used the 2bk>0bk contrast (along with two other contrasts) to define MD regions (Assem et al., 2020), reproduced in **Figure 1a**. We have previously identified that the strongest and most domain-general MD activations overlap with 10 HCP MMP1.0 areas that we have labelled core MD (Assem et al., 2024, Assem et al., 2020). Lateral frontal core MD regions are surrounded by white borders in **Figure 1a**.

For an initial overview of category-specific responses, we examined group average activation maps for each category contrasted against the average of the remaining 3 categories (averaging across difficulty levels i.e. 0bk and 2bk). In striking contrast to the MD activation pattern, each category map highlights a patchwork of focal activations mainly outside of core MD areas, highlighting their distinct spatial organization (**Figure 2a**).

**Figure 2.**
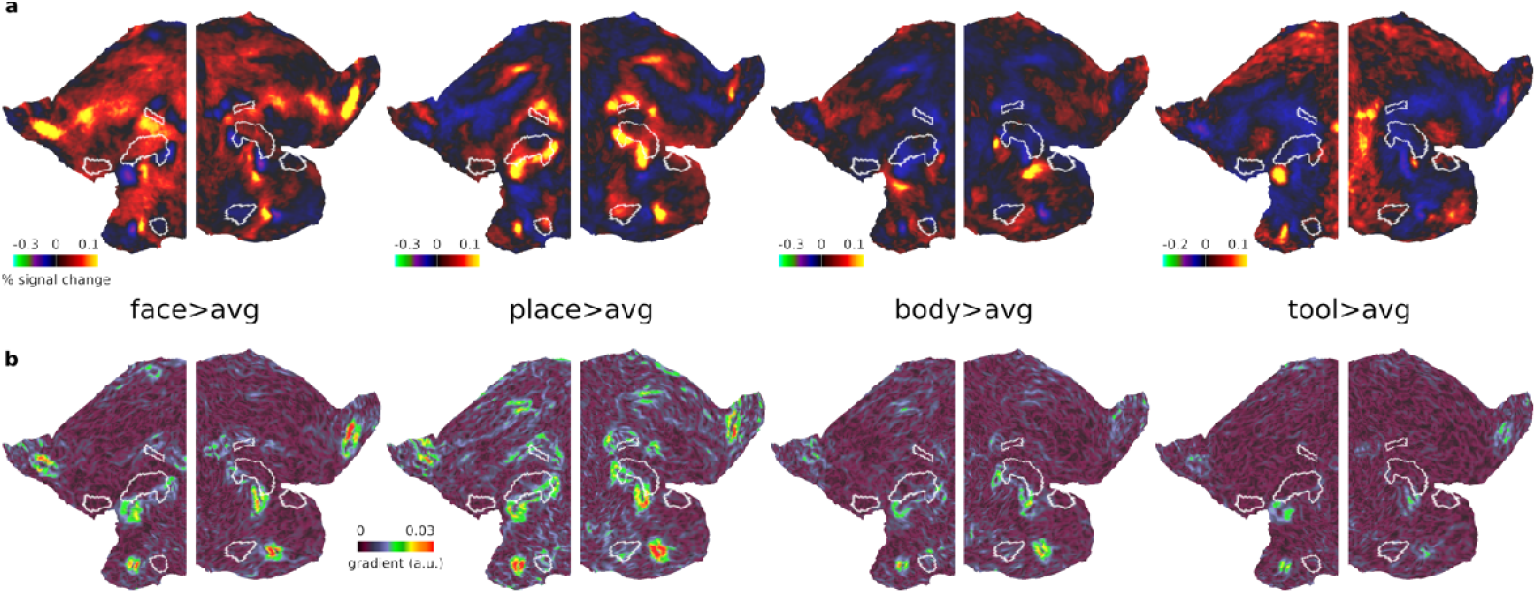
Unthresholded HCP activation and gradient maps. (a) Bilateral flat cortical maps displaying category-biased activations (e.g. face>average of the three other categories) averaging across 0bk and 2bk conditions. **(b)** Gradient maps for each contrast. Data available at https://balsa.wustl.edu/3xnK7.

To quantitatively delineate these category-biased activations near MD regions, we performed a vertex-level statistical analysis within a predefined bilateral frontal cortical mask that encompassed lateral core MD and their immediately adjacent areas (**Figure 1b, see Methods**). To identify significant vertices, we performed a one sample t-test (*p*<0.05 FDR corrected) for each category compared to each of the others (e.g. face>place, face>body, face>tool). Then we performed a conjunction analysis across the three contrasts to identify the commonly significant vertices. From these, we selected the top 5% activated vertices to delineate clustered vertices, or patches. This analysis identified multiple category-biased patches across both hemispheres (**Figure 1c**; see **Supplementary Figure 1** for their relation to the labelled HCP MMP1.0 atlas). We highlight three notable locations. In the insular region, face and place-biased patches are located anterior to core MD region AVI. In the ventral PFC region, we observed an interdigitated pattern of all four categories along a strip ventral and posterior to core MD areas, especially prominent in the left hemisphere ventral to core MD area p9-46v. Although all patches were located just outside core MD areas, one place patch partially overlaps with core MD area IFJp. In the dorsal frontal region, we identify at least 3 place patches surrounding core MD area i6-8, and at least 2 face patches inferior to it, though their exact locality and extent varied between hemispheres.

To assess the replicability of these findings, we split the dataset into two independent halves [P210 and V210 groups, the parcellation and validation groups used in (Glasser et al., 2016)] and repeated the same analysis. Indeed, the split-half analysis confirmed most of the identified patches and their close proximity to core MD regions, demonstrating robust spatial consistency across subsets of participants (**Figure 1d**).

Finally, to refine the assessment of the spatial relationship between these category-biased patches and core MD areas, we computed gradient maps (using subject-specific vertex areas, **see Methods**) for each of the category group average maps in **Figure 2b**. A gradient map is the first spatial derivative of the activation map and will not highlight where category preferences are but will indicate regions where preferences change rapidly relative to neighboring vertices. Hence, gradients can be in similar locations across the different category maps. Importantly, this gradient map analysis avoids the reliance on arbitrary statistical thresholds used to define clusters, providing a more continuous and unbiased representation of how activation preferences change across the cortex. **Figure 2c** indeed highlights prominent gradients, which reflect rapid shifts in category preferences, consistently located at the boundaries of core MD regions.

Collectively, these findings underscore the adjacency of category-biased patches to core MD regions, suggesting a finely tuned spatial organization within the lateral frontal cortex. In the next section, we explore if these findings replicate in an independent dataset with a diverse range of tasks.

### Consistency of category-biased patches across independent datasets and tasks

To investigate the generality of these findings, we analyzed an independent HCP-style dataset (n=37) in which participants performed 4 distinct tasks: 3 block-designed executive function (EF) tasks (n-back, switching and stop signal), and one event-related temporal sequence task, where participants had to memorize the order of the presented images (**see Methods**). All tasks used the same face and place stimuli, allowing us to identify activation preferences for these two categories.

We first examined the 3 executive tasks, each containing easy and hard blocks. For the main analysis, we averaged across the 3 EF tasks for each participant. As reported in Assem et al 2024, activations for the hard>easy EF contrasts strongly co-activate core MD (reproduced in **Figure 3a**) reinforcing their domain-generality beyond the HCP tasks. In contrast, the group average face>place contrast (averaged across tasks and difficulties), revealed focal category-biased activations, closely mirroring the patterns observed in the HCP dataset (**Figure 3b**).

**Figure 3.**
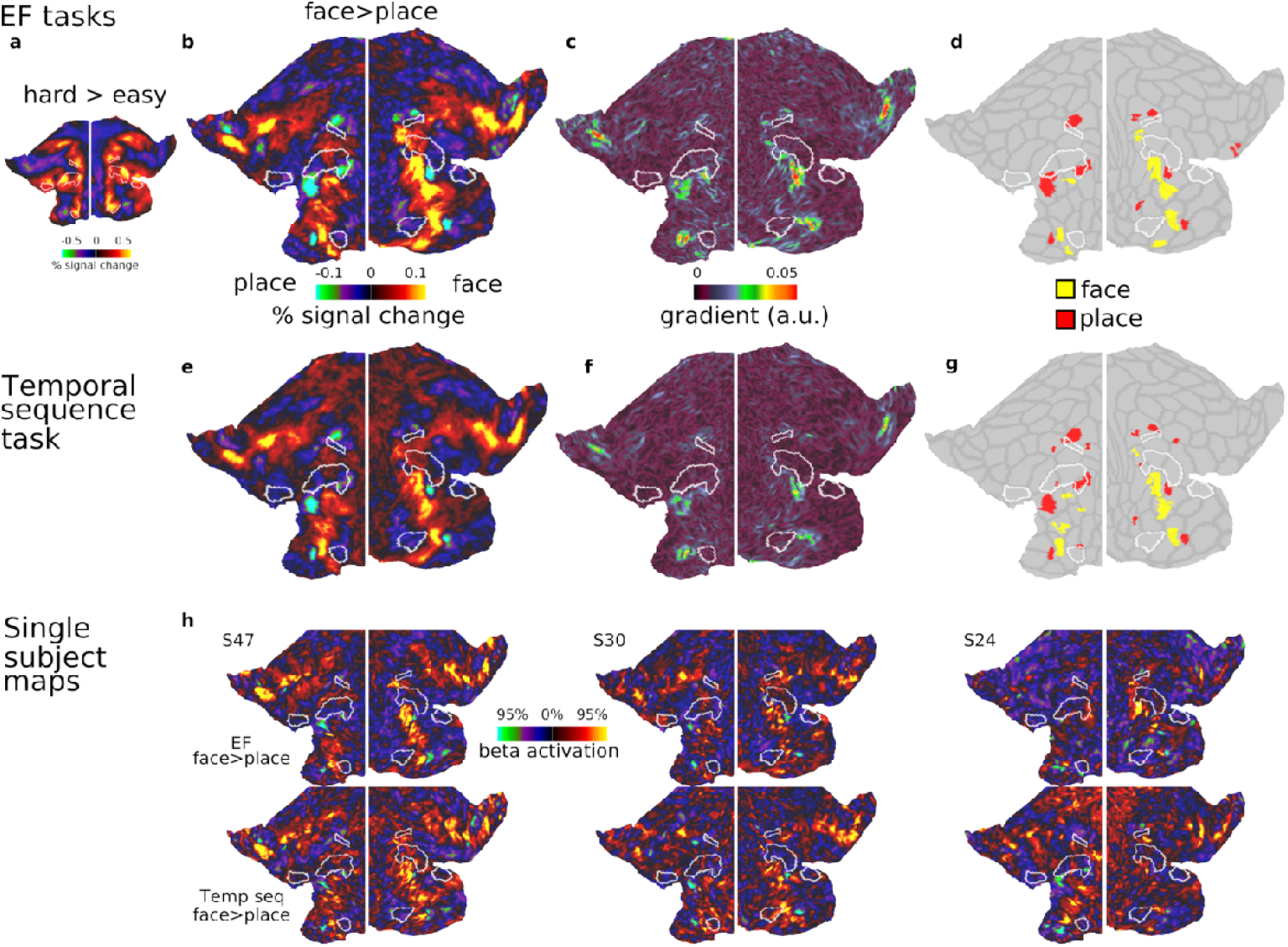
Category selective patches in the CBU dataset. (a) Flat map displaying group average activation map for the hard>easy contrast, averaged across three executive tasks (n-back, switch, stop-signal) **(b)** Group average activation map for the face>place contrast, averaged across tasks and difficulty levels. Focal category-biased patches are located adjacent to core MD regions. **(c)** Gradient map for the face>place contrast. **(d)** Statistically defined category-biased patches (face>place). **(e)** Group average activation map for the face>place contrast during the temporal sequence task, revealing a strikingly similar topography to the executive map in (b). **(f)** Gradient map for the face>place contrast in the temporal sequence task. **(g)** Statistically defined category-biased patches from the temporal sequence task **(h)** Three single subject examples of face>place activation maps from the executive task (top) and temporal sequence task (bottom), demonstrating consistency of category-biased patches at the individual level. Data in this figure and all single subject maps are available at https://balsa.wustl.edu/z8rN3.

Using the statistical cluster identification method from the previous section (i.e. one sample-test on the contrast of face>place), we identified most of the same face and place patches in the previous section (**Figure 3d**). These included the patches in the insular region, along with the ventral and dorsal PFC regions. Notable exceptions were the absence of two face patches in the dorsal aspect of the PFC bilaterally, which were apparent in the group average map but did not survive statistical thresholds-likely due to the smaller sample size and task-functional preferences. The place patch overlapping with IFJp was only identified in the left hemisphere, again suggesting task-specific variations. Importantly, this analysis robustly highlights the consistent locations of face and place patches adjacent to core MD regions. A gradient map of the face>place contrast (**Figure 3c**) emphasized that sharp changes in category preferences are adjacent to core MD regions.We next analyzed the temporal sequence task, performed in a separate session by the same participants. The group average of face>place contrast revealed a topography strikingly similar to that observed in the executive tasks (**Figure 3e**). Applying the same cluster identification method identified many of the same patches as the EF tasks, with minor differences, likely attributable to statistical thresholding (**Figure 3g**). The gradient map (**Figure 3f**) similarly highlights prominent shifts sparing core MD areas, consistent with the patterns observed in the EF tasks.

To further illustrate the consistency of the face and place patches at the individual level, we include example single-subject activation maps from EF and temporal sequence tasks (**Figure 3h**). These maps demonstrate consistent category-biased patterns adjacent to core MD borders, reinforcing the reliability of these findings.

Together, these results demonstrate a consistent spatial organization of category-biased patches adjacent to core MD regions across independent datasets and diverse task paradigms. In the next section, we explore their functional preferences through ROI analysis.

### Cross-validating the functional preferences of category-biased and core MD areas

In the previous section we demonstrated the robustness of category-biased patches encircling core MD regions in the lateral frontal cortex across two independent datasets. Here, we aim to cross-validate their functional responses. Specifically, we used patches defined in one dataset to estimate responses in the other, focusing on the face>place and hard>easy contrasts. This analysis tests whether category-biased areas, like sensory-biased regions, exhibit consistent domain-specific responses and whether their activity scales with task difficulty.

First, we used all the face and place patches defined in the HCP dataset, as well as the lateral core MD patches, and examined the average responses of each ROI to EF tasks. For this analysis, we averaged activation values across all vertices within each patch, and then across all patches of the same type (i.e. face, place and MD) separately for each subject. For the face>place contrast (averaged across easy and hard conditions), face patches had stronger activations for the face condition, while place patches showed the opposite pattern (**Figure 4a**; one sample t-test averaging across hemispheres, face ROIs *t*_*36*_*=*9.7, place ROIs *t*_*36*_*=*5.6, *face vs place t*_*36*_*=*9.5, all *p*s<0.0001). In contrast, lateral frontal core MD areas showed no statistically significant category preferences (**Figure 4a**; t_36_*=*0.6, *p*=0.52). Next, we analyzed the hard>easy contrast (averaged across face and place categories). Here, place patches showed increased activations, while face patches showed decreased activations (**Figure 4b**; face ROIs *t*_*36*_*=*5.3, place ROIs *t*_*36*_*=*18.8, all *p*s<0.0001). As expected, core MD areas also showed a difficulty response, that was stronger than the response in place patches (MD ROIs *t*_*36*_*=*28.0, place vs MD ROIs *t*_*36*_*=*7.9, *p*<0.0001). These results are consistent with sensory-biased regions (Assem et al., 2022), indicating that, like MD regions, some domain-specific areas also show a task difficulty effect.

**Figure 4.**
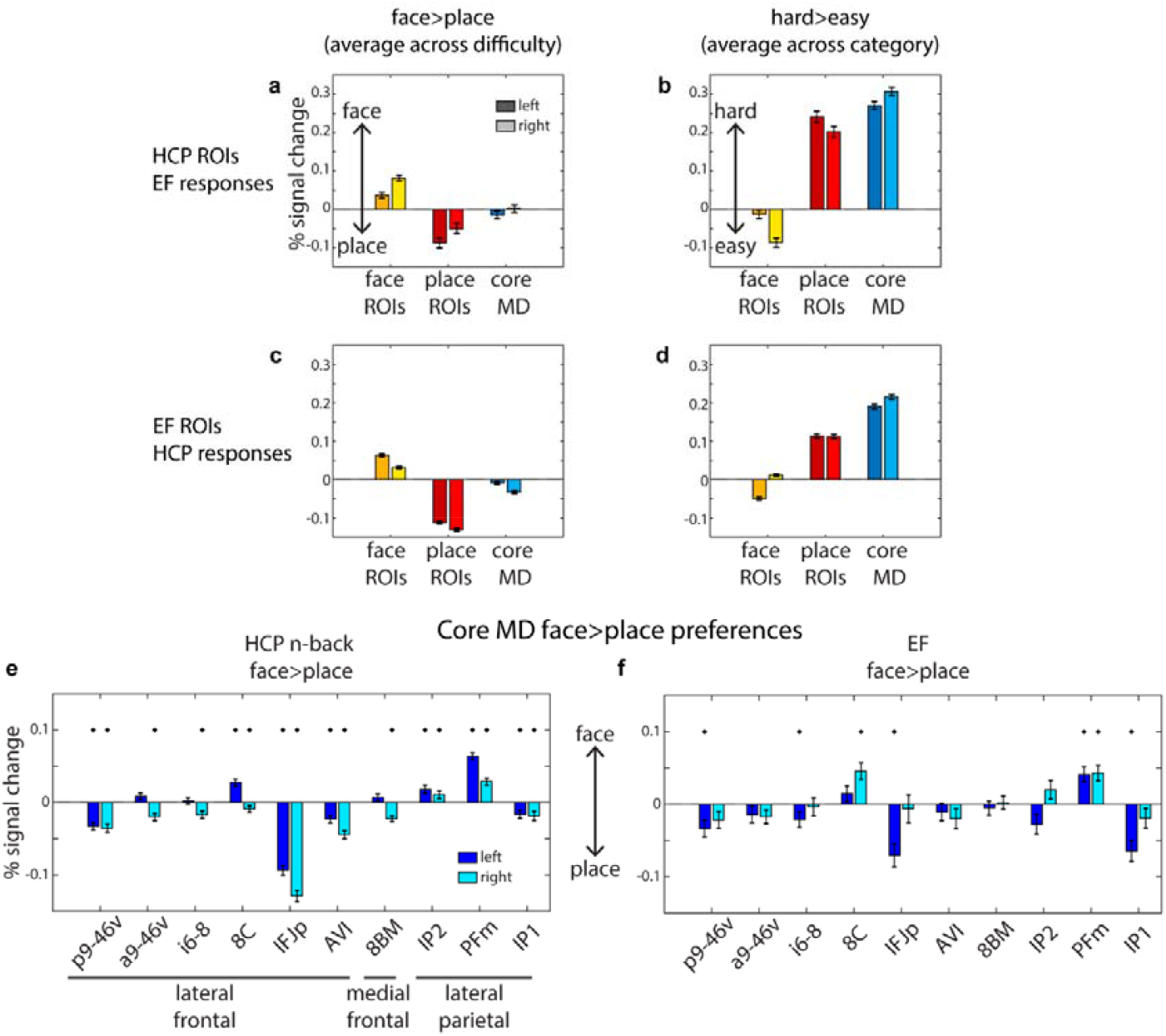
Functional profiles of category-biased and core MD regions. (a, b) Estimated response for the EF tasks to the HCP’s n-back defined face and place ROIs. Core MD regions are based on the definition in Assem et al 2020. **(c, d)** Estimated responses for the HCP n-back task to the EF defined face and place ROIs. **(e, f)** Individual core MD areal category preferences during the HCP n-back task **(e)** and the EF task **(f)**. In all panels, error bars are standard error of means. * p<0.05 uncorrected one-sample t-test.

To investigate if the face and place patches are modulated by EF task type, we repeated the face>place contrast analysis for each EF task separately (**Supplementary Figure 2**). Most patches were sensitive to all tasks though with some functional preferences. The n-back task activated 8 of 9 face patches and 5 of 12 place patches bilaterally (one-sample t-test, *p*<0.05 Bonferroni corrected within each patch for 3 tasks). The switch task activated 7 of 9 face and all 12 place patches. The stop task activated 4 of 9 face and 8 of 12 place patches. These results confirm broad consistency of category preferences across tasks, though with some possibility of differential recruitment.

We repeated the same analysis using EF-defined face and place patches to estimate their responses during the HCP n-back task. For the face>place contrast (averaged across difficulty), we replicated strong category preferences for each ROI, in the expected directions (**Figure 4c**; face ROIs *t*_*448*_*=*16.5, place ROIs *t*_*448*_*=32*.*0*, face vs place *t*_*448*_*=34*.*9, p*<0.0001). Core MD areas displayed minimal place preferences (*t*_*448*_*=*5.6, *p*<0.0001), with place ROIs showing approximately 6 times stronger place responses than core MD preferences (mean MD ROIs response = 0.02, mean place ROIs response = 0.12, *t*_*448*_=29.7, *p*<0.0001). For the hard>easy contrast, we again observed stronger activations for the place and MD areas but not the face areas (**Figure 4d**; face ROIs *t*_*448*_*=*16.5, place ROIs *t*_*448*_*=32*.*0*, MD ROIs *t*_*448*_*=34*.*9, p*<0.0001).

Both cross-validated analysis revealed that place-biases are sensitive to task difficulty, whereas face biases are not. To examine whether this reflects differences in behavioral difficulty, we compared task performance across categories. In the HCP task, there was no significant difference between place (90.8±9.0%) and face (91.0±8.9%) stimuli (paired sample t-test; face vs place t_445_=0.52 *p=*0.6). Body parts were the most difficult (82.6±11.0%; paired sample t-test vs each of the other categories *p*<0.001, Bonferroni corrected for 12 pairwise comparisons) followed by the tools category (87.2±10.0%). In the CBU executive dataset, face stimuli (94.6±3.3%) were only slightly easier than place stimuli (93.1%±2.99%; t_36_=3.37 *p*=0.002). Given closely matched performance, it is unlikely that, in place patches, selective response to place tasks was heavily driven by task difficulty.

Finally, we probed category preferences of all cortical core MD areas individually, including those outside lateral frontal cortex. Given that previous findings found a general visual bias in core MD responses (Assem et al 2022), we explored whether there might be heterogeneity in their category-preferences. Using the face>place contrast (averaged across difficulty), we consistently found across both datasets that a subset of parietal core MD regions (IP2 and PFm) and frontal area 8C exhibited face preferences, whereas all remaining ones showed place preferences (**Figure 4e-f**). Taken together, these results demonstrate consistent category preferences across datasets for the face and place patches and reveals subtle heterogeneity in category preferences within core MD areas.

### Sensory-biased preferences of the category-biased patches

To further contextualize the category-biased patches, we compared them with previously reported sensory-biased regions in the frontal lobe (Assem et al., 2022; Michalka et al., 2015; Noyce et al., 2022; Tobyne et al., 2017). In the same CBU dataset, participants also performed an auditory n-back task, which we previously used to localize sensory-biased patches by contrasting visual and auditory n-back tasks (Assem et al., 2022). This overlap in tasks and participants provides a unique opportunity to directly examine the spatial relationship between category-biased and sensory-biased patches within individuals.

For a visual overview, we overlaid the EF-defined face and place patches over the group average activations from the same participants for the visual vs auditory n-back contrast (averaged across difficulty; **Figure 5a;** Assem et al., 2022). This revealed substantial overlap between the place patches and visually biased activations. In contrast, face patches show less consistent overlap, with some appearing sandwiched between visual and auditory patches - most notably in the ventral frontal face area of the left hemisphere (see inset of **Figure 5a**).

**Figure 5.**
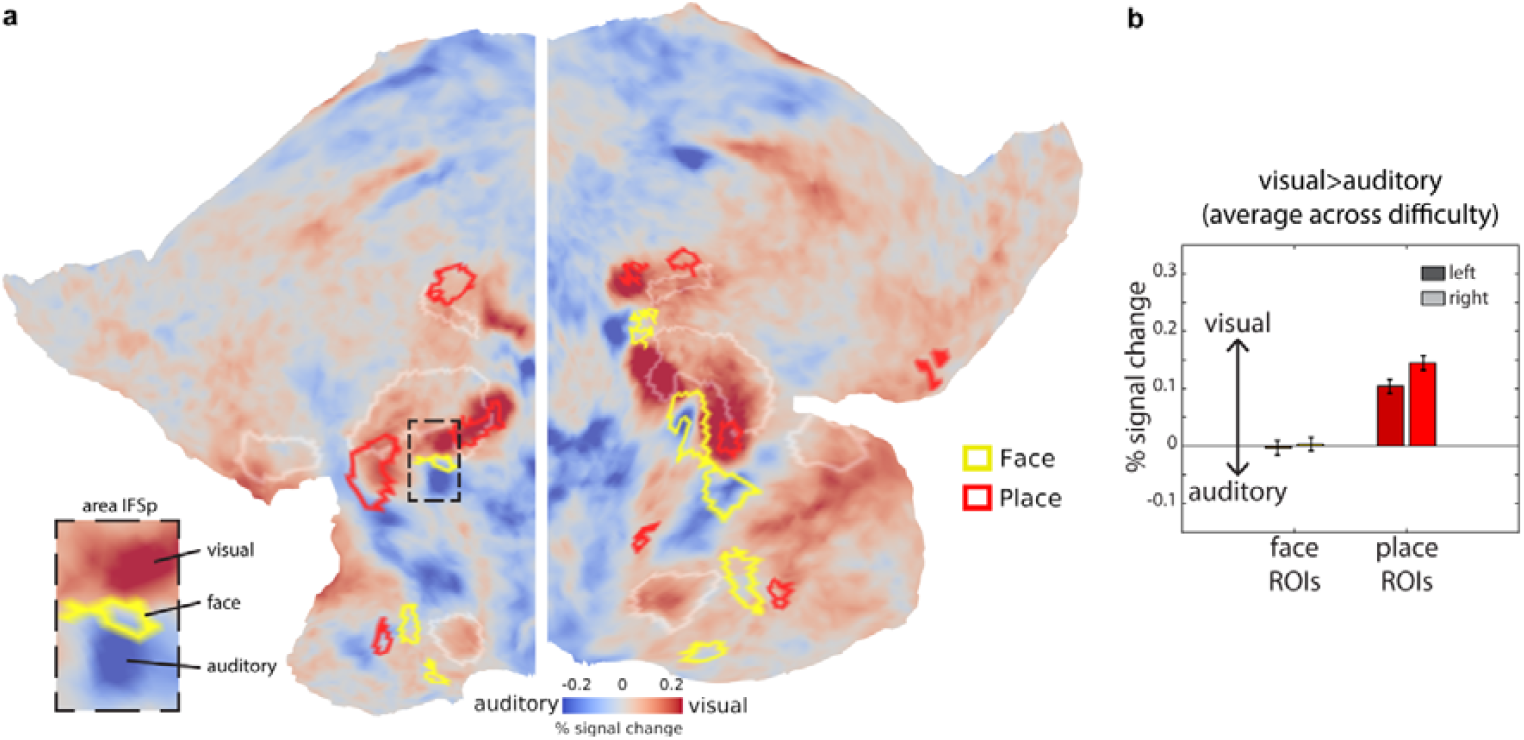
Visual and auditory preferences of category-biased regions. (a) Flat map displaying group average activation map for the visual>auditory n-back contrast (averaged across difficulty) for the same CBU EF participants. Red borders are place-biased patches and yellow borders are face-biased patches, both defined from the CBU EF dataset (**Figure 3d**). Inset shows enlarged patch around left area IFSp in the ventral frontal cortex (see **Supplementary Figure 1** for HCP MMP1.0 areal labels and borders). Note the face patch is sandwiched between two opposing peaks of visual and auditory preferences. **(b)** Estimated responses for the EF visual vs auditory n-back contrast to the EF defined face and place ROIs. Data in this figure are available at https://balsa.wustl.edu/VDgr6

To quantify these observations, we extracted responses from EF-defined face and place patches during the visual vs auditory n-back contrast (**Figure 5b**). This analysis confirmed a strong visual bias for the place patches (one sample t-test, t_36_=11.3 *p*<0.001) and no significant sensory preferences for the face patches (t_36_=0.05 *p*=0.96; face vs place interaction t_36_=8.4 *p*<0.001).

This findings reveal an interesting link beween visual and place biases, suggestting they might share inherent functional properties. Further, the positioning of some face patches between visual and auditory biases gives further evidence for fine-grained functional interdigitation around MD regions.

## Discussion

The fine-grained functional organization of the lateral PFC remains a key challenge in understanding how this region supports complex cognition. In this study, we demonstrated, across independent datasets and cognitive tasks, that the human lateral PFC consists of multiple interdigitated category-biased patches responsive to faces, places, tools and body parts. Importantly, most of these patches consistently spared the domain-general core MD regions, encircling them and lying in close adjacency (**Figures 1-3**). These findings point to a highly structured and fine-grained organization within the PFC, where interdigitated domain-specific and domain-general patches may represent a fundamental organizational principle. We previously proposed that this arrangement facilitates flexible cognitive control by enabling the integration of task-specific signals from functionally segregated systems into core MD regions (Assem et al., 2022, 2020; Duncan et al., 2020). The presence of multiple patches for the same category suggests that these patches are embedded within different brain networks either large scale e.g. default mode, dorsal attention (Ji et al., 2019) or more fine-grained (Rajimehr et al., 2024). This widespread distribution would allow category-biased patches to access distinct types of information (e.g. memory-vs action-related) relevant to each category. MD regions, in turn, provide the workspace where diverse and context-specific information can intersect. In this framework, each category-biased patch contributes specialized information to spatially adjacent MD regions, creating the control signals tailored to the demands of the current task. This model predicts that different tasks, even when using identical stimuli of the same category, should differentially engage these patches. This prediction was supported by preferential engagement of the category-biased patches across the three EF tasks (**Supplementary Figure 2**).

We observed a particularly rich concentration of category-biased patches along a ventral strip of the PFC (**Figures 1 and 3**). Interestingly, previous work on sensory-biased areas (Assem et al., 2022; Michalka et al., 2015; Noyce et al., 2017) identified visual and auditory biased patches along the same strip (**Figure 5**). Leveraging the CBU dataset, where the same participants performed both sensory and category tasks, we found that visually biased patches overlapped with place patches, whereas face patches did not show a dominant sensory preference. Functionally, place-biased patches exhibited similar functional properties to visual-biased patches, both co-activating with MD regions in response to task difficulty (Assem et al., 2022). Notably, the difficulty sensitivity of place patches is unlikely to be explained by behavioral difficulty alone, as participants found both face and place tasks similarly challenging. In contrast, face biases did not align with auditory biases and were positioned between auditory and visual biased regions, suggesting a higher, more integrative role – potentially linking to social or theory of mind networks (Koster-Hale and Saxe, 2011; Paunov et al., 2019; Rajimehr et al., 2024). Their insensitivity to task difficulty further aligns with broader evidence showing that social brain regions are generally less modulated by cognitive demand (Diachek et al., 2020; Koster-Hale and Saxe, 2011; Paunov et al., 2019).

The HCP dataset was particularly rich, allowing us to investigate four categories. Notably, tool-biased responses were generally located farther from core MD regions, closer to premotor cortex, suggesting close links to motor actions (**Figures 1 & 2**). As a result, only a few tool-biased vertices survived our constrained masks adjacent to core MD. However, one particularly interesting region ventral to the core MD area p9-46v showed an unusually high concentration of all four categories. In this region, tool biases appeared largely overshadowed by stronger place and body biases (**Figures 1 & 2**).

Though not presented here, we note that, in these datasets, category biases are also visible in other parts of the association cortices, including medial PFC and medial and lateral parietal regions. Some of these biases have been documented in previous studies (Silson et al., 2019; Steel et al., 2025). Here, we focused on the lateral PFC because of its unique organizational feature of densely packing representations-i.e. smaller distances between functional areas compared to other cortical territories (Xu et al., 2022). However, systematic examination of category biases beyond lateral PFC remains an important direction for future studies.The diversity of category-biased patches raises the possibility that even more such regions exist. For example, a recent study suggest response biases to geometric shapes near the intra-parietal sulcus (Sablé-Meyer et al., 2024). Higher-resolution studies, including a broader range of categories, may reveal an even finer mosaic of category-biased patches within the PFC and across the association cortex.

This study adds to growing evidence that the PFC is much more functionally heterogenous than captured in classic lesion-based characterizations of large frontal territories (Assem et al., 2024; Braga and Buckner, 2017; Glasser et al., 2016). Domain-general and domain-specific regions are extensively interdigitated through out the PFC, challenging traditional views that distinct broad PFC regions support separate cognitive control processes. We propose that through this distributed overlapping architecture, the PFC can dynamically integrate multiple strems of information, supporting the construction of flexible cognitive control models that guide adaptive behaviour (Assem et al., 2024; Duncan, 2025).

## Data availability

Data used for generating each of the imaging-based figures are available on the BALSA database (https://balsa.wustl.edu/study/mljn6). Selecting the URL at the end of each figure will link to a BALSA page that allows downloading of a scene file plus associated data files; opening the scene file in Connectome Workbench will recapitulate the exact configuration of data and annotations as displayed in the figure.

## Acknowledgments

We would like to thank Reza Rajimehr and the Woolgar lab for discussions and comments on this work.

## Supplementary Figures

**Supplementary Figure 1.**
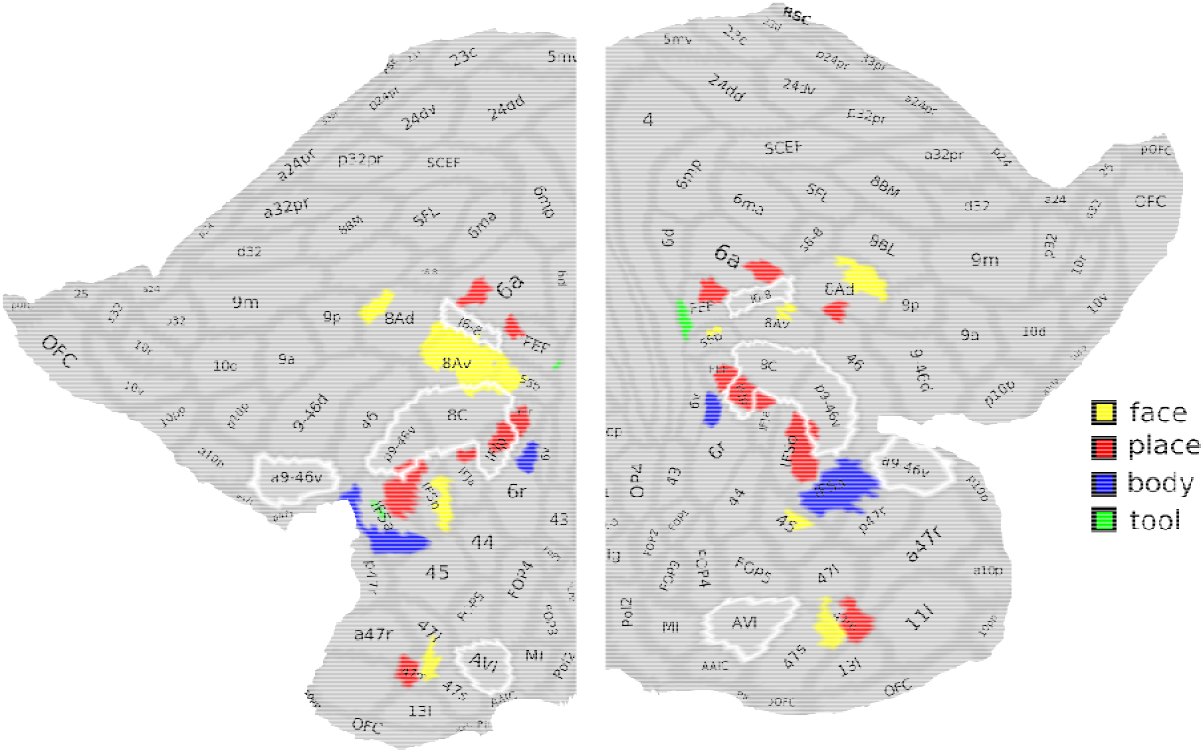
Category-biased patches in the HCP dataset from **Figure 1** overlaid on the areal labels of the HCP MMP1.0 (Glasser et al., 2016). Data in this figure are available at https://balsa.wustl.edu/3q0lx

**Supplementary Figure 2.**
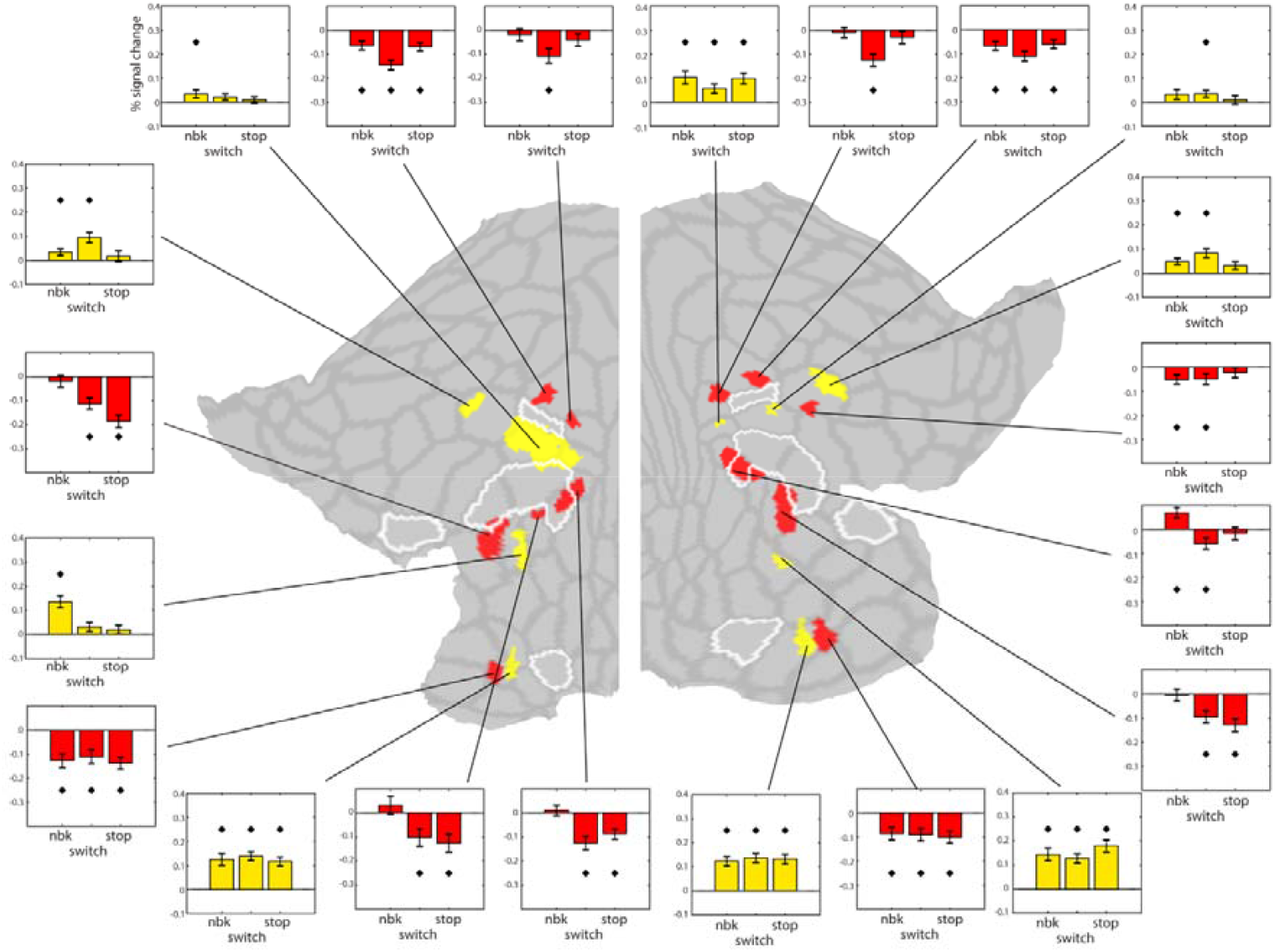
Estimated response (face minus place) for each EF task [n-back (nbk), switch, stop] to the HCP’s n-back defined face and place ROIs. Core MD regions (white borders) are based on the definition in (Assem et al., 2020). * indicates p<0.05 Bonferroni corrected within each patch for 3 tasks.

